# The *C. elegans* RIG-I homolog *drh-1* mediates the Intracellular Pathogen Response upon viral infection

**DOI:** 10.1101/707141

**Authors:** Jessica N. Sowa, Hongbing Jiang, Lakshmi Somasundaram, Guorong Xu, David Wang, Emily R. Troemel

## Abstract

Mammalian RIG-I-like receptors detect viral dsRNA and 5’ triphosphorylated RNA to activate transcription of interferon genes and promote antiviral defense. The *C. elegans* RIG-I-like receptor DRH-1 promotes defense through antiviral RNA interference, but less is known about its role in regulating transcription. Here we describe a role for *drh-1* in directing a transcriptional response in *C. elegans* called the Intracellular Pathogen Response (IPR), which is associated with increased pathogen resistance. The IPR includes a set of genes induced by diverse stimuli including intracellular infection and proteotoxic stress. Previous work suggested that the proteotoxic stress caused by intracellular infections might be the common trigger of the IPR, but here we demonstrate that different stimuli act through distinct pathways. Specifically, we demonstrate that DRH-1/RIG-I is required for inducing the IPR in response to Orsay virus infection, but not in response to other triggers like microsporidian infection or proteotoxic stress. Furthermore, *drh-1* appears to be acting independently of its known role in RNAi. Interestingly, expression of the replication competent Orsay virus RNA1 segment alone is sufficient to induce most of the IPR genes in a manner dependent on RNA dependent RNA polymerase activity and on *drh-1*. Altogether, these results suggest that DRH-1 is a pattern-recognition receptor that detects viral replication products to activate the IPR stress/immune program in *C. elegans*.

**Importance:** *C. elegans* lacks homologs of most mammalian pattern recognition receptors, and how nematodes detect pathogens is poorly understood. We show that the *C. elegans* RIG-I homolog *drh-1* mediates induction of the Intracellular Pathogen Response (IPR), a novel transcriptional defense program, in response to infection by the natural *C. elegans* viral pathogen Orsay virus. *drh-1* appears to act as a pattern-recognition receptor to induce the IPR transcriptional defense program by sensing the products of viral RNA-dependent RNA polymerase activity. Interestingly, this signaling role of *drh-1* is separable from its previously known role in antiviral RNAi. In addition, we show that there are multiple host pathways for inducing the IPR, shedding light on the regulation of this novel transcriptional immune response.

## Introduction

RIG-I-like receptors (RLRs) are an ancient family of cytoplasmic pattern recognition receptors that detect viral dsRNA and 5’ triphosphorylated RNA to trigger antiviral immune responses (Ahmad and Hur, 2015; Lässig and Hopfner, 2017). In mammals this family includes RIG-I and MDA5, which contain a common N-terminal tandem caspase activation and recruitment domain (2CARD), a central DExD/H box motif helicase domain, and a zinc-binding C-terminal domain (CTD) (Yoneyama et al., 2005). After the helicase and CTD bind to viral RNAs, these receptors trigger a signaling cascade via interaction of the CARD domain with the mitochondrial activator of virus signaling (MAVS) protein, which ultimately results in the activation of the IRF3 and NF-κB transcription factors (Wu and Hur, 2015). These transcription factors trigger downstream defense gene expression, including an IRF3-mediated antiviral type-I interferon response (Gebhardt et al., 2017). Given the importance of RLRs in antiviral defense and autoimmunity, further understanding their evolution and signaling mechanisms in different contexts could provide new avenues for treatment of viral infections and autoimmune diseases.

RIG-I is one of the few pattern recognition receptors conserved between mammals and the model nematode *C. elegans*. Notably, *C. elegans* lacks the cytoplasmic pattern recognition receptors cGAS-STING and NLR proteins, and also lacks canonical Toll-like receptor/ NF-κB signaling (Cohen and Troemel, 2015; Irazoqui et al., 2010; Pujol et al., 2001; Pukkila-Worley, 2016). Furthermore, *C. elegans* lacks obvious homologs of IRF3 and MAVS as well as interferon ligands and receptors. However, *C. elegans* does possess three genes that encode RIG-I-like receptor homologs: dicer-related helicases 1, 2, and 3 *(drh-1, drh-2*, and *drh-3*) (Duchaine et al., 2006; Tabara et al., 2002). Like RIG-I and MDA5, these genes encode helicase and CTD domains, but have a divergent N terminal domain. DRH-1 was originally identified in *C. elegans* as a protein that interacts with the dsRNA-binding protein RDE-4 and Dicer/DCR-1 to process dsRNA into siRNAs (Tabara et al., 2002). Subsequently, *drh-1* was found to mediate an antiviral response to several types of viral infection in *C. elegans* (Ashe et al., 2013; Coffman et al., 2017; Gammon et al., 2017; Guo et al., 2013; Lu et al., 2009).

Functional analysis indicated that the helicase domain from human RIG-I could substitute for the helicase domain in *C. elegans drh-1* to promote anti-viral defense against both of these infections (Guo et al., 2013). Interestingly, a small deletion in the CTD of *drh-1* was shown to underlie natural variation in *C. elegans* strains susceptible to infection by the Orsay virus (Ashe et al., 2013). The Orsay virus has a positive sense ssRNA genome composed of just two segments, the RNA1 segment which contains an open reading frame (ORF) encoding an RNA-dependent RNA polymerase (RDRP), and the RNA2 segment which contains an ORF encoding a capsid protein and an ORF implicated in viral exit (Félix et al., 2011; Yuan et al., 2018).

Because *C. elegans* lacks interferon signaling, characterization of the antiviral mechanism of *drh-1* has focused on its role in mediating antiviral RNA interference. The role for *drh-1* in triggering a transcriptional response to viral infection has been less clear (Tanguy et al., 2017).

Intriguingly, *C. elegans* responds to Orsay virus infection by upregulating mRNA expression of a set of genes that are also induced during infection with pathogens in the microsporidia phylum (Bakowski et al., 2014; Chen et al., 2017; Reddy et al., 2019). Like viruses, microsporidia are obligate intracellular pathogens that are natural pathogens of the *C. elegans* intestine (Troemel et al., 2008). However, microsporidia are molecularly distinct, as they are eukaryotic pathogens in the fungal kingdom. We have named the common transcriptional response to microsporidia and the Orsay virus the Intracellular Pathogen Response (IPR), as it appears to represent a novel stress/immune pathway including many genes upregulated in the intestine (Bakowski et al., 2014; Reddy et al., 2017). The IPR is regulated by two members of the *pals* gene family, which is named for a protein signature of unknown biochemical function, and has a single member each in mouse and human (Leyva-Díaz et al., 2017; Reddy et al., 2017). *C. elegans* mutants defective in the gene *pals-22* constitutively express IPR genes, and have increased resistance to virus and microsporidia infection, which is reversed when increased IPR gene expression is suppressed by mutations in *pals-25* (Reddy et al., 2019). In addition to increased pathogen resistance, *pals-22* mutants have several other phenotypes including increased thermotolerance dependent on ubiquitin ligase components, increased RNAi interference, increased susceptibility to the extracellular bacterial pathogen *Pseudomonas aeruginosa*, slowed development, and shortened lifespan (Leyva-Díaz et al., 2017; Reddy et al., 2017). Thus, activation of IPR genes is associated with a broad rewiring of *C. elegans* physiology.

How are IPR genes induced? Given that they can be induced not just by infection, but also by proteotoxic stressors like prolonged heat stress and proteasome blockade, a simple hypothesis was that intracellular infections by microsporidia and virus were creating proteotoxic stress that led to IPR gene induction (Bakowski et al., 2014). However, here we show that there are additional layers of complexity in the regulation of IPR induction. Specifically, we show that induction of IPR gene expression by viral infection requires the *drh-1* receptor. *drh-1* mutants are defective in inducing IPR genes in response to viral infection but not in response to other IPR triggers, including microsporidian infection and proteotoxic stress. Furthermore, we show that activation of this transcriptional response does not require known DRH-1 interacting proteins RDE-4 or DCR-1, and thus appears to occur via a mechanism that is distinct from the antiviral RNAi pathway. Finally, we use RNA-seq to show that the IPR transcriptional program can be induced by ectopic expression of the Orsay virus RNA1 segment, and this induction depends on RDRP activity and on *drh-1*. Together these results suggest that *drh-1* acts a pattern recognition receptor in *C. elegans* to sense viral replication products and activate the IPR stress/immune program.

## Results

### The RIG-I ortholog *drh-1* is required for induction of the Intracellular Pathogen Response by Orsay virus infection, but not by other stressors

Infection of *C. elegans* by the Orsay virus or by the microsporidian species *N. parisii* induces a common set of genes as part of the IPR, including the gene *pals-5*. While the *pals-5* gene is of unknown function, the *pals-5p∷GFP* reporter provides a convenient read-out for the IPR (Bakowski et al., 2014). To investigate the role of *drh-1* in inducing the IPR, we examined *pals-5p∷GFP* expression in the background of a *drh-1(ok3495)* partial deletion allele. We found that, in contrast to wild-type animals, Orsay virus infection did not induce *pals-5p∷GFP* reporter expression in *drh-1(ok3495)* mutants, despite these mutants carrying a higher viral load as assessed by qRT-PCR of the viral RNA1 genome segment (Figure 1a-c). Because the *drh-1(ok3495)* allele has only a partial deletion, we used CRISPR/Cas9 editing to generate an additional deletion allele of *drh-1*, in which the entire genomic locus was removed (Figure 1d). This complete deletion allele, *drh-1(jy110)*, also failed to upregulate the IPR reporter when infected with Orsay virus (Figure 1e) and carried a higher viral load compared to wild-type (*WT*) worms (Figure 1f). For simplicity, from this point onward the *drh-1(ok3495)* allele will be denoted as *drh-1(−)*.

**Figure 1:**
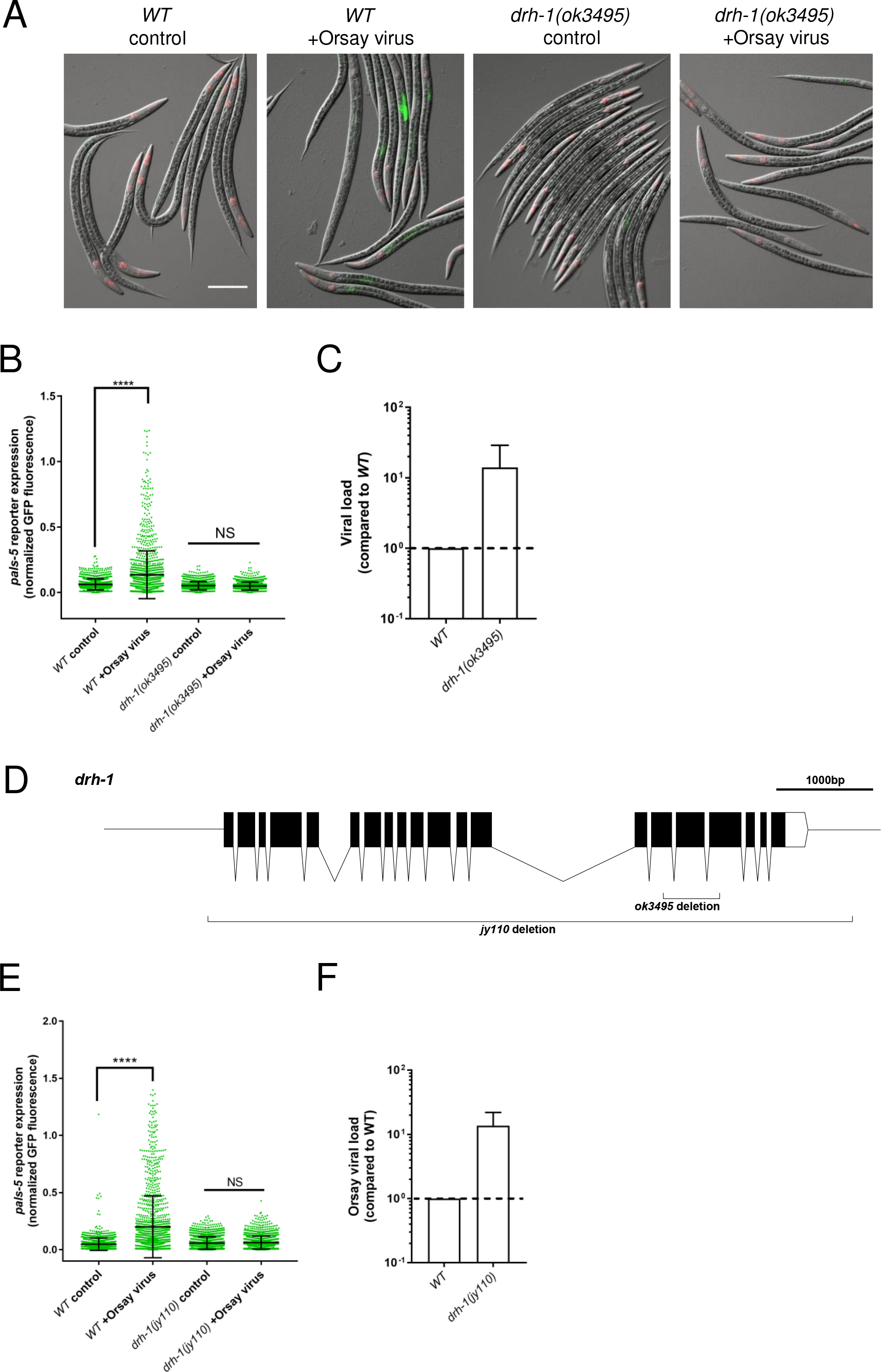
*drh-1* is required for induction of the *pals-5p∷GFP* IPR reporter upon viral infection. **A)** *pals-5p∷GFP* IPR reporter fluorescence after Orsay virus infection in *WT* vs *drh-1(ok3495)* backgrounds. Worms were infected with Orsay virus as L1 larvae and imaged at 24 hpi. Scale bar = 100 μm. **B)** *pals-5p∷GFP* IPR reporter fluorescence after Orsay virus infection in *WT* vs *drh-1(ok3495)* backgrounds. Worms were infected with Orsay virus as L2 larvae and reporter expression assayed at 24 hpi, each dot represents an individual animal. Graph shows one representative experiment out of three independent experimental replicates. **C)** qRT-PCR analysis of Orsay viral load of samples from Figure 1b. Graph shows combined results of two independent experimental replicates. Error bars represent standard deviation (SD). **D)** Genomic location of *jy110* and *ok3495* deletions in *drh-1*. **E)** *pals-5p∷GFP* IPR reporter fluorescence after Orsay virus infection in *WT* vs *drh-1(jy110)* backgrounds. Worms were infected with Orsay virus as L2 larvae and reporter expression assayed at 24 hpi, each dot represents an individual animal. Graph shows a representative experiment out of three independent experimental replicates. **F)** qRT-PCR analysis of Orsay viral load of samples from Figure 1e. Graph shows combined results of three independent experimental replicates. Error bars represent standard deviation (SD). Panels B, and E: Green fluorescence and length of animals were measured for >100 individual animals using a COPAS Biosorter. Means were compared using One-way ANOVA with Bonferroni correction. NS = Not significant, ****p<0.0001

Interestingly, we found no defect in IPR reporter activation by *N. parisii* infection or proteasome blockade in either of the *drh-1* mutant alleles, and we did not observe a substantial difference in *N. parisii* pathogen load in *drh-1* mutants compared to *WT* controls (Figure 2). Indeed, in both *drh-1* deletion alleles proteasome blockade or *N. parisii* infection resulted in stronger GFP reporter activation than was seen in *WT* controls, with the effect stronger in the *drh-1(jy110)* complete deletion background (Figure 2a-f). We also saw robust activation by prolonged heat stress in both *drh-1* mutant alleles (Figure 2g-h). Furthermore, the increased *pals-5p∷GFP* expression seen in a *pals-22* mutant background was not compromised by a mutation in *drh-1* (Figure 2i), indicating that this pathway for IPR induction is not dependent on *drh-1*. Thus, we found that *drh-1* is required specifically for *pals-5p∷GFP* expression in response to viral infection, but not for response to other triggers of *pals-5p∷GFP* expression.

**Figure 2:**
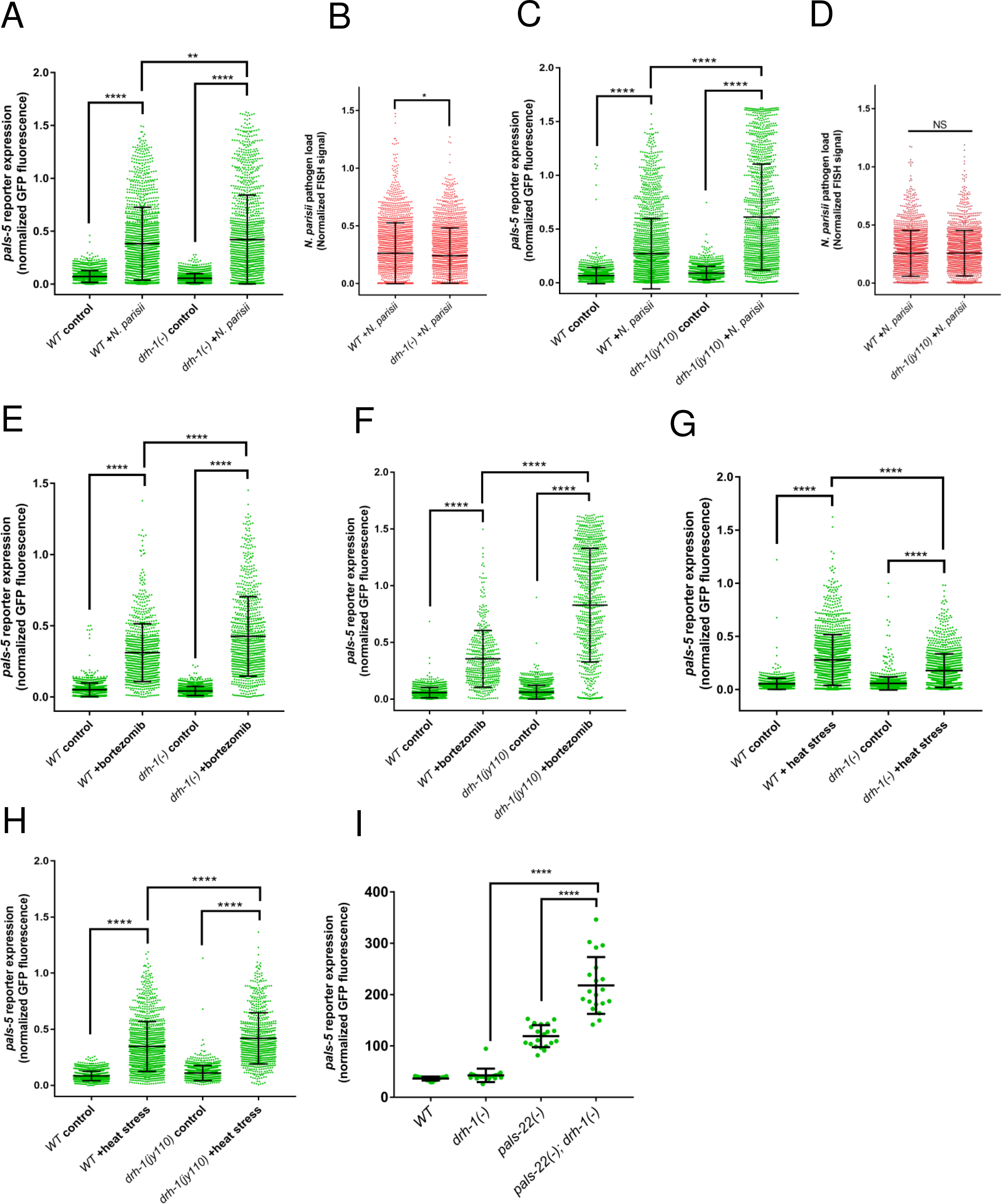
*drh-1* is not required for induction of the *pals-5p∷GFP* IPR reporter upon non-viral triggers. **A** and **C)** *pals-5p∷GFP* reporter fluorescence during *N. parisii* infection. Animals were infected with *N. parisii* spores as L1 larvae and reporter expression measured at 30 hpi. Graph shows a representative experiment out of three (Panel A) or two (Panel C) independent experimental replicates. **B** and **D)** *N. parisii* pathogen load assayed by *N. parisii* specific FISH staining. Graph shows a representative experiment out of three (Panel B) or two (Panel D) independent experimental replicates. **E** and **F)** *pals-5p∷GFP* IPR reporter expression in worms treated with 2.5 μM bortezomib. Graph shows a representative experiment out of two (Panel E) or three (Panel F) replicates. **G** and **H)** *pals-5p∷GFP* reporter expression in worms subjected to 24 h of 28°C heat stress beginning at L4 stage. Graphs shows representative experiments out of three independent experimental replicates for each. **I)** Quantification of intestinal *pals-5p∷GFP* IPR reporter fluorescence in *pals-22(jy1)* single mutants and *pals-22(jy1); drh-1(ok3495)* double mutants. Graph shows quantification from one experiment (n=20 animals for each strain). For panels A - H, fluorescence and length of animal was measured for >100 individual worms using a COPAS Biosorter, with each animal represented as a dot. Means were compared using One-way ANOVA with Bonferroni correction. NS = Not significant, *p<0.05, **p<0.01, ***p<0.001, ****p<0.0001.

### *drh-1* acts independently of known RNAi factors to induce IPR mRNA expression in response to Orsay virus infection

To obtain a broader picture of the requirement for *drh-1* in IPR gene expression, we performed qRT-PCR analysis for a panel of IPR genes including those of unknown biochemical function *pals-5* and *F26F2.1*, mRNA decapping enzyme *eol-1*, and ubiquitin ligase complex component *skr-5* (Bakowski et al., 2014; Reddy et al., 2017, 2019). As a negative control we included *skr-1*, which is a ubiquitin ligase complex component that is not induced upon infection (Bakowski et al., 2014; Reddy et al., 2017). We compared expression levels in Orsay infected worms vs mock infected controls at 12 hours and 24 hours post-infection (hpi) in *WT* and *drh-1(−)* mutant backgrounds (Figure 3). We observed that at 12 hpi there was a >10-fold induction of *pals-5*, *F26F2.1*, and *eol-1* in the Orsay-infected *WT* worms, but no detectable induction in the *drh-1(−)* background (Figure 3a) despite the fact that they were carrying a much higher viral load than their *WT* counterparts (Figure 3b). We observed a similar pattern at 24 hpi, with *pals-5, F26F2.1*, and *eol-1* being induced >100 fold by Orsay infection in the *WT* background, while much less induction was seen in the *drh-1(−)* background despite *drh-1(−)* carrying an increased viral load (Figure 3c-d). No induction of *skr-1* was observed in *WT* or *drh-1(−)* at 12 hpi or 24 hpi (Figure 3a, c). In contrast to the lack of IPR induction in *drh-1* mutants at 12 hpi, some induction of *pals-5, F26F2.1* and *eol-1* was observed in the *drh-1* mutant background at 24 hpi, raising the possibility that there could be a *drh-1*-independent induction of IPR genes by Orsay virus that functions at later stages of infection or with higher viral loads. Of note, at 24 hpi *skr-5* showed higher expression in the infected *drh-1* mutants compared to infected *WT* (Figure 3c) demonstrating that unlike the other IPR genes assayed, *skr-5* induction does not require *drh-1*. Analysis of the *drh-1(jy110)* full deletion allele at 24 hpi confirmed these findings (Figure 3e-f). Together, these data demonstrate that *drh-1* is required to induce several IPR genes in response to Orsay virus infection.

**Figure 3:**
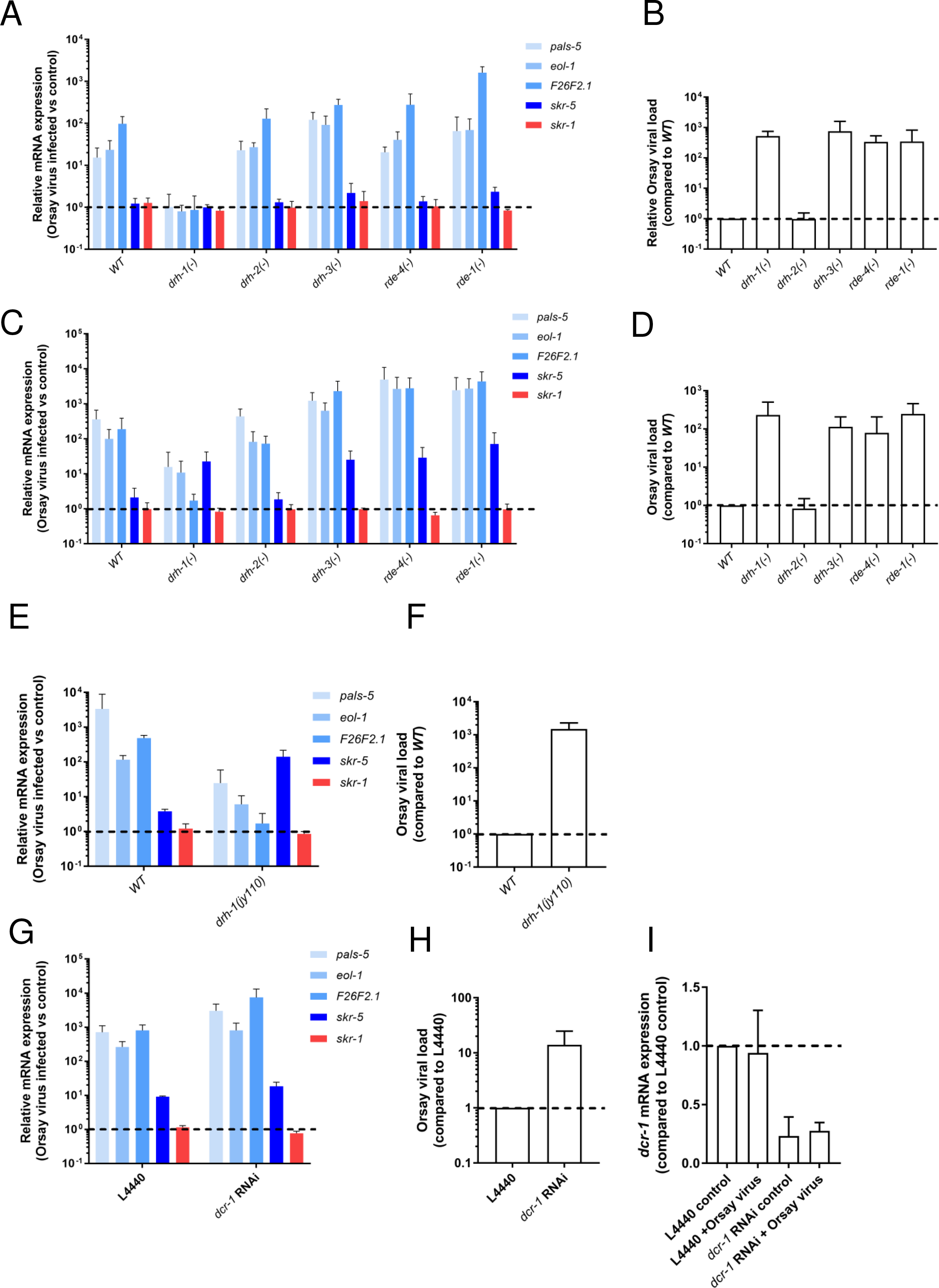
DRH-1 acts independently of other RNAi factors to mediate IPR induction. **A)** Induction of IPR genes analyzed by qRT-PCR in RNAi pathway mutants infected with Orsay virus at 12 hpi. **B)** qRT-PCR assay of Orsay viral load in RNAi pathway mutants at 12 hpi of Orsay virus infection. **C)** Induction of IPR genes analyzed by qRT-PCR in RNAi pathway mutants at 24 hpi of Orsay virus infection. **D)** qRT-PCR assay of Orsay viral load in RNAi pathway mutants at 24 hpi of Orsay virus infection. **E)** Induction of IPR genes by qRT-PCR in *drh-1(jy110)* mutants infected with Orsay virus at 24 hpi. **F)** qRT-PCR assay of Orsay viral load in *drh-1(jy110)* mutants at 24 hpi of Orsay virus infection. **G) I**nduction of IPR genes by qRT-PCR in worms treated with RNAi against *dcr-1* after 24 hpi of Orsay virus infection. **H)** qRT-PCR assay of Orsay virus pathogen load in worms treated with RNAi against *dcr-1* at 24 hpi of Orsay virus infection. **I)** qRT-PCR assay of *dcr-1* transcript levels in samples from panel G. *dcr-1* levels expressed as ratio to levels in L4440 mock infected control. All panels: Graphs show average of three independent experimental replicates, error bars represent SD. Panels A, C, E, and G: IPR gene expression is shown as a ratio between Orsay infected vs mock-infected controls for each genotype. Panels B, D, F, and H: Orsay viral load as assessed by Orsay RNA1 transcript levels compared to levels in Orsay-infected *WT*.

We next investigated whether other components of the antiviral RNAi pathway play a role in the virus-induced activation of IPR gene expression. In addition to *drh-1*, the *C. elegans* genome encodes two other DExD/H box RNA helicases with homology to mammalian RLRs, *drh-2* and *drh-3* (Duchaine et al., 2006; Tabara et al., 2002). DRH-2 is not thought to have a direct role in antiviral RNAi, but DRH-3 participates in the production of secondary siRNAs (Guo et al., 2013; Lu et al., 2009). Other factors involved in the production of siRNAs targeting viral transcripts include RDE-4 and DCR-1, which both interact directly with DRH-1, potentially as part of the complex that binds viral dsRNAs to initiate cleavage, and RDE-1, an argonaute that binds primary siRNAs (Duchaine et al., 2006; Guo et al., 2019; Parrish and Fire, 2001; Tabara et al., 2002). To test the role of these genes we infected *rde-1, rde-4, drh-2*, and *drh-3* mutants with Orsay virus for 12 and 24 hpi (infections were performed in parallel with *WT* and *drh-1(−)* infections described above). qRT-PCR analysis for a panel of IPR genes showed that none of these other antiviral RNAi mutants had impaired upregulation of IPR gene expression in response to viral infection (Figure 3a, c). At both 12 and 24 hpi IPR gene expression levels were higher in *rde-1, rde-4*, and *drh-3* mutants compared to *WT*, which is consistent with the higher viral load observed in these mutants (Figure 3b, d). Importantly, *drh-1* mutants attained a similar viral load to *rde-1, rde-4*, and *drh-3* mutants at both time points, while displaying drastically less induction of IPR genes. (Figure 3a-d). No defects in IPR activation were observed in *drh-2* mutants, and they showed comparable viral loads to *WT* worms at both time points (Figure 3a-d).

Because null mutations in the *C. elegans* Dicer homolog *dcr-1* lead to sterility or lethality (Billi et al., 2014), we used RNAi knockdown to determine whether *dcr-1* was required for IPR activation by Orsay virus. Worms raised on *dcr-1* RNAi beginning at L1 stage and infected with Orsay virus at L4 showed higher levels of IPR activation at 24 hpi than worms raised on RNAi vector control and also carried a higher viral load (Figure 3g-i), indicating that *dcr-1* is not required for IPR activation by Orsay virus. Altogether, these results indicate that *drh-1*, but not other canonical RNAi factors like *dcr-1*, are required for mediating induction of IPR gene expression upon viral infection.

### IPR gene expression is induced in a manner dependent on *drh-1* and on the activity of viral RNA-dependent RNA polymerase

Previous work indicated that transgenic expression and replication of only Orsay virus RNA1, the viral genome segment containing the RDRP, was sufficient to activate the *pals-5p∷GFP* IPR reporter, and that this activation was lost when a mutation was introduced that ablated the polymerase activity of the RDRP (and therefore also ablated replication of the RNA1 segment) (Jiang et al., 2017). Here we investigated whether this effect was dependent on *drh-1*. First, we compared IPR reporter expression in transgenic animals expressing Orsay RNA1(wt) or Orsay RNA1 D601A(RDRP defective mutant) under the control of a heat-shock promoter in a *WT* or *drh-1* mutant background. We found that, in agreement with previous work, heat-shock induced expression and replication of the Orsay virus RNA1(wt) in the *WT* background led to an increase in IPR reporter expression, whereas ectopic expression of the RNA1(mt) did not (Figure 4a). Importantly, we found that in the *drh-1* mutant background, neither the RNA1(wt) nor the RNA1(mt) construct was able to induce IPR reporter expression, indicating that *pals-5p∷GFP* induction by Orsay virus RNA1(wt) is dependent on *drh-1* (Figure 4a). We tested the specificity of the requirement for *drh-1* in activating the IPR in response to RNA1 RDRP activity by crossing the RNA1(wt) array into the *rde-1, rde-4*, and *drh-3* mutant backgrounds. We found that neither *rde-1, rde-4*, nor *drh-3* were required to induce *pals-5p∷GFP* expression in response to heat-shock induced ORV RNA1(wt) expression (Figure 4b), confirming that IPR activation resulting from RNA1 RDRP activity occurs via a pathway distinct from the canonical antiviral RNAi pathway.

**Figure 4:**
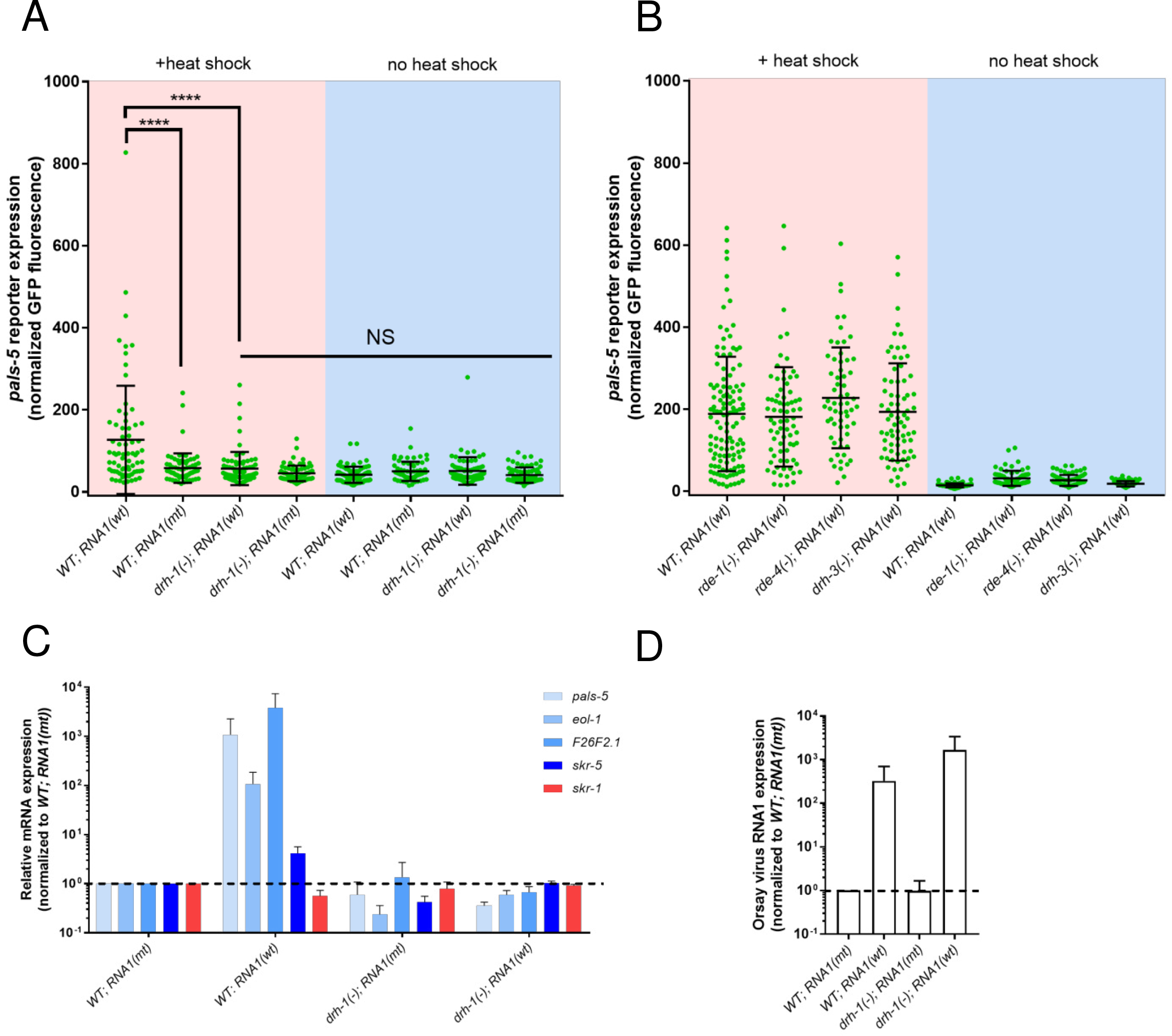
RDRP activity of Orsay RNA1 is required for DRH-1 mediated IPR activation. **A)** Quantification of *pals-5p∷GFP* reporter fluorescence after heat shock induction of RNA1(wt) or RNA1(mt) expression in *WT* and *drh-1(ok3495)* background. Graph shows combined results of 3 independent experimental replicates (n≥70). Means were compared using One-way ANOVA with Bonferroni correction. NS = Not significant, ****p<0.0001 **B)** Quantification of *pals-5p∷GFP* reporter fluorescence after heat shock induction of RNA1(wt) expression in *WT*, *rde-1(ne219), rde-4(ne301)*, and *drh-3(ne4253)* backgrounds. Graph shows combined results of 3 independent biological replicates for each mutant genotype (n≥60) and 5 replicates for *WT* (n=125). **C)** qRT-PCR analysis of IPR gene expression after heat shock induction of RNA1(wt) vs RNA1(mt) in the *WT* and *drh-1(ok3495)* backgrounds. IPR gene expression is compared to expression in *WT; RNA1(mt)*. **D)** qRT-PCR analysis of RNA1 transcript levels after heat shock induction of RNA1(wt) and RNA1(mt) in *WT* and *drh-1(ok3495)* backgrounds. RNA1 transcript levels are compared to RNA1 levels in *WT; RNA1(mt)*. Panels C-D: Graphs show combined results of 3 independent experimental replicates, error bars represent SD.

We next investigated whether other IPR genes are induced by expression of RNA1, and whether this induction is dependent on *drh-1*. We performed qRT-PCR on animals expressing RNA1(wt) and RNA1(mt) in both *WT* and *drh-1(−)* backgrounds. First, we found that ectopic expression of RNA1(wt) induced all IPR genes tested, and this induction was dependent on *drh-1* (Figure 4c). Analysis of the Orsay RNA1 transcript levels showed that RNA1 accumulated to higher levels in the *drh-1* mutants compared to *WT* controls (Figure 4d), indicating that the lack of induction was not due to lack of this RNA1 trigger.

### The full repertoire of IPR genes can be induced by Orsay virus RNA-dependent RNA polymerase activity, dependent on *drh-1*

Given that ectopic expression of RNA1(wt) was sufficient to induce expression of a subset of IPR genes (Figure 4c), we examined the full transcriptomic response to RNA1(wt) expression using RNA-seq, and investigated whether responses were dependent on *drh-1*. To minimize the contribution of the RNAi pathway, we used the *rde-1(ne219)* RNAi-deficient mutant background for all of our comparisons. Specifically, we compared these strains: 1) *rde-1(−); RNA1(wt)*, 2) *rde-1(−); RNA1(mt)*, 3) *drh-1(−); rde-1(−); RNA1(wt)*, and 4) *drh-1(−); rde-1(−); RNA1(mt)*, and used heat shock to induce RNA1 expression in all strains before harvesting RNA for RNA-seq (Figure 5a). For simplicity, we refer to these strains as 1) *drh-1(+)* animals expressing RNA1(wt), 2) *drh-1(+)* animals expressing RNA1(mt), 3) *drh-1(−)* mutants expressing RNA1(wt), and 4) *drh-1(−)* mutants expressing RNA1(mt).

**Figure 5:**
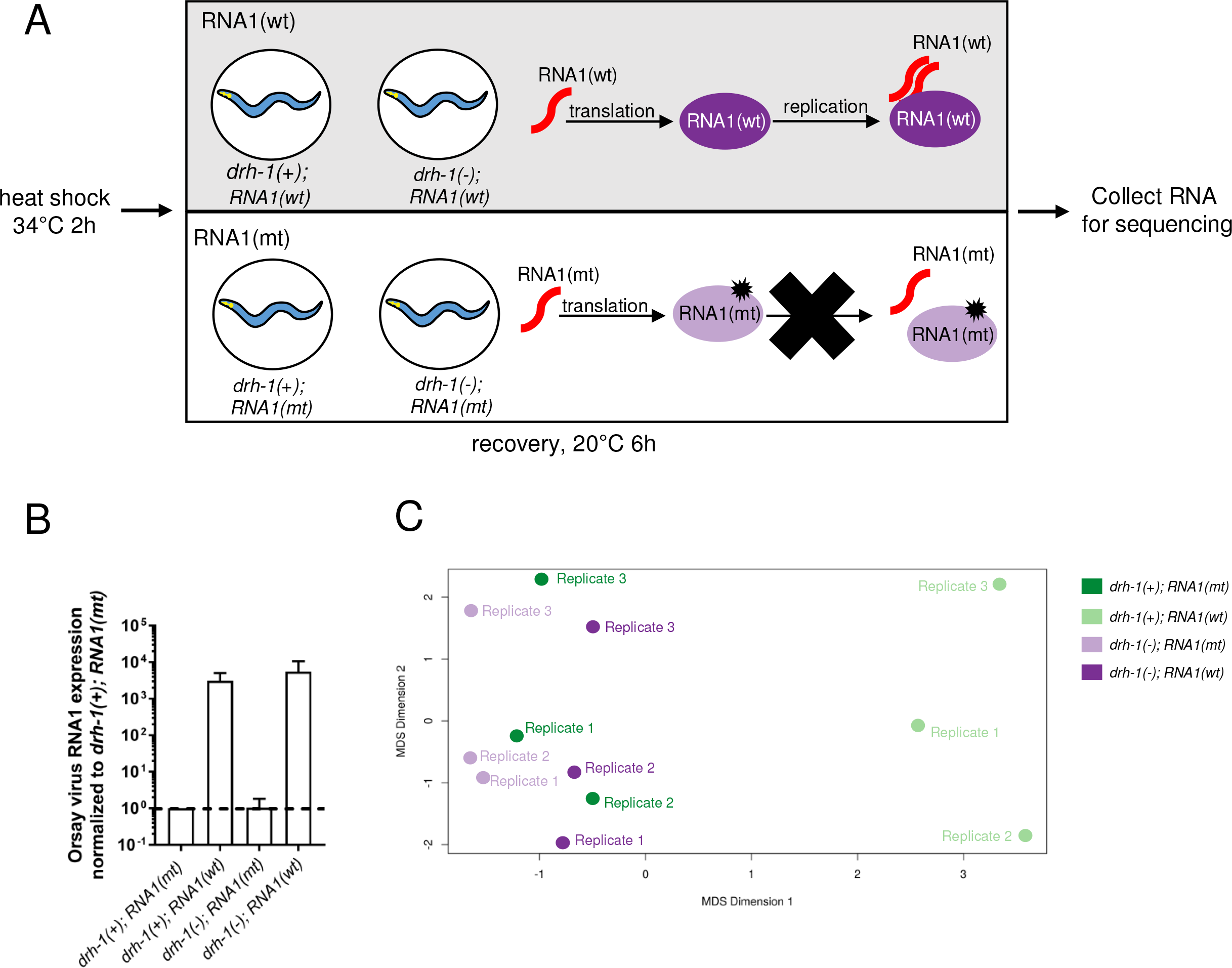
Characterization of the transcriptional response to Orsay RNA1. **A)** Schematic of sample preparation for RNA-seq. Synchronized populations of L1 larvae for all four genotypes (*drh-1(+); RNA1(mt), drh-1(+); RNA1(wt), drh-*1(-); *RNA1(mt)*, and *drh-1(−); RNA1(wt))* were obtained by COPAS Biosort-based isolation of animals with the transgenic array marker *myo-2p∷YFP* to obtain a relatively pure population of transgene-positive animals. After sorting, worms were plated with food and incubated at 20°C for 2 days until the L4 stage. At the L4 stage, worms were subjected to 2h heat shock at 34°C to induce expression of RNA1. Worms were recovered at 20°C for 6h, then harvested for RNA isolation. **B)** qRT-PCR analysis of RNA1 transcript levels in samples used for RNA-seq. RNA1 transcript levels are compared to RNA1 levels in *drh-1(+); RNA1(mt)*. Graph shows combined results for all three RNA-seq replicates, bars represent mean and error bars represent SD. **C)** MDS plot of all three RNA-seq replicates.

First, we used qRT-PCR to measure the level of RNA1 expressed in the triplicate RNA samples used for RNA-seq analysis of these four strains. Here we saw that the levels of RNA1 were much higher in animals expressing RNA1(wt) compared to RNA1(mt) (Figure 5b), which is expected given that RNA1 is a polymerase that amplifies the copy number of RNA1. RNA1 expression levels were slightly higher on average in the *drh-1(−)* animals expressing RNA1(wt) compared to the *drh-1(+)* animals expressing RNA1(wt) (Figure 5b). Then we performed RNA-seq, and used multidimensional scaling analysis on the RNA-seq results to determine which strain had the most distinct gene expression profile. Here we found that mRNA expression in *drh-1(+)* animals expressing RNA1(wt) diverged most dramatically from mRNA expression in the other three strains (Figure 5c), indicating a transcriptional response dependent on RDRP activity and wild-type *drh-1*.

When comparing *drh-1(+)* animals expressing RNA1(wt) to *drh-1(+)* animals expressing RNA1(mt), we found 194 genes significantly upregulated and 1 significantly downregulated (Figure 6a, Supplemental Table 1). Of these, only 26 genes were also significantly upregulated in *drh-1(−)* animals expressing RNA1(wt) vs. *drh-1(−)* animals expressing RNA1(mt) (Figure 6a-b, Supplemental Table 2). These results indicate that the transcriptional response to RDRP activity is largely dependent on *drh-1*. Importantly, there was also a large degree of overlap between genes induced by RDRP activity and “canonical” IPR genes, which were defined by being induced by *N. parisii* infection and regulated by *pals-22/25* (Reddy et al., 2019). A majority of the IPR genes induced by RNA1(wt) were *drh-1* regulated (Figure 6b, Supplemental table 2). For example, predicted ubiquitin ligase components like Cullin *cul-6*, which is required for increased thermotolerance in *pals-22* mutants (Reddy et al., 2017), as well as the Skp-related protein *skr-4* and F-box protein *fbxa-75*, were induced by RNA1(wt) expression in a *drh-1*-dependent manner (Supplementary Table 2). We also compared the genes induced by RNA1 activity with a previously published data set of genes induced in the *rde-1* mutant background by Orsay virus infection (Chen et al., 2017), and found that the majority of the RDRP activity-induced genes were also induced during natural Orsay infection (Figure 6a-c, Supplemental Table 2). While there were genes listed as virally induced that were not significantly upregulated by expression of RNA1(wt), most of these were induced but failed to meet the significance cut-off (Supplemental Table 3). Therefore, the IPR, which is a common response to molecularly divergent pathogens like microsporidia and virus, can be induced by expression of replication-competent Orsay RNA1 in a manner dependent on RDRP activity and *drh-1*.

**Figure 6:**
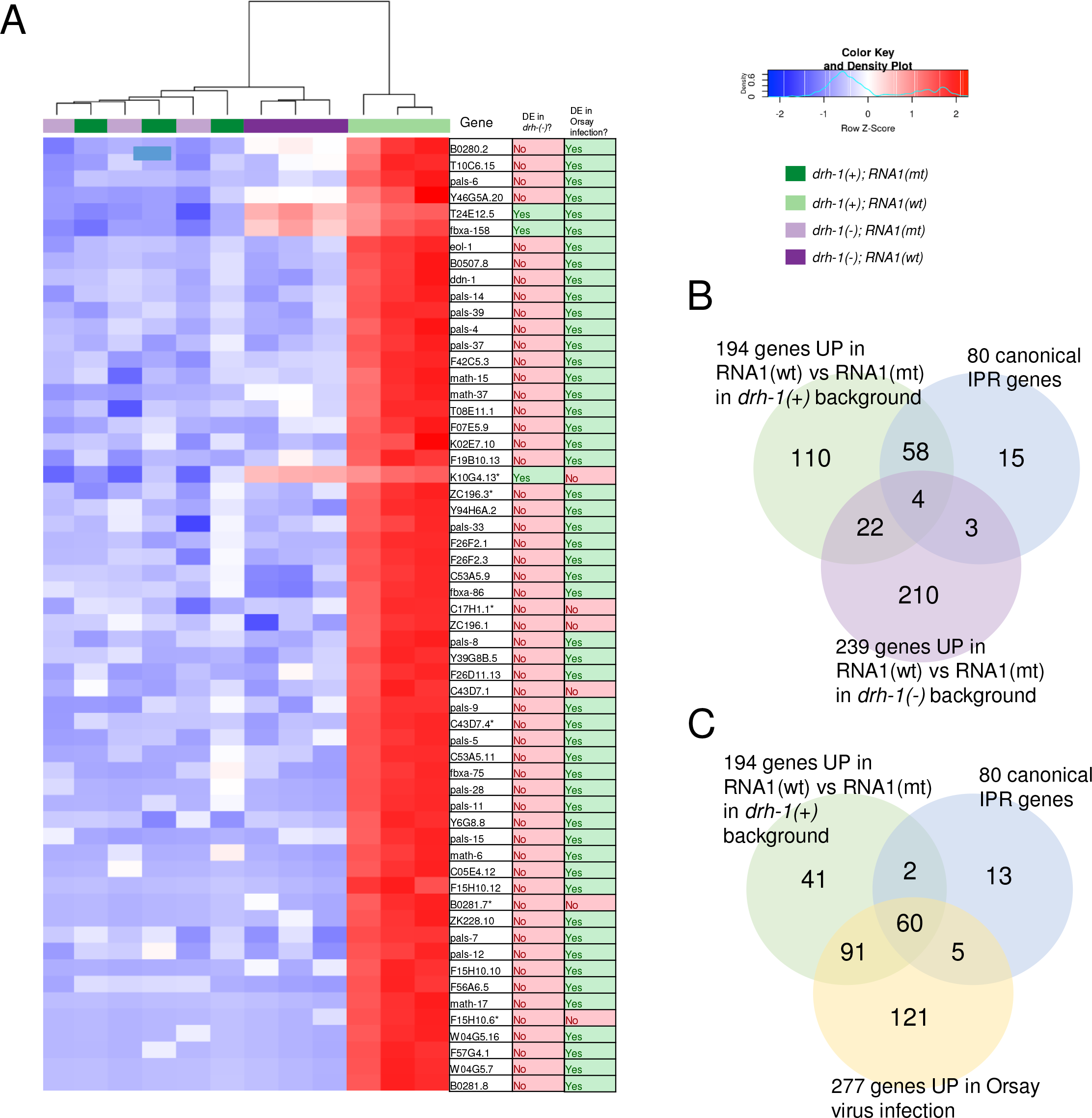
Ectopic expression of Orsay RNA1 induces IPR gene repertoire in an RDRP-dependent and *drh-1*-dependent manner. **A)** Heat map showing top (p<0.003) differentially expressed genes in *drh-1(+); RNA1(wt)* vs *drh-1(+); RNA1(mt)*. (*) denotes pseudogenes. Columns indicate whether gene was also differentially expressed in *drh-1(−); RNA1(wt)* vs *drh-1(−); RNA1(mt)*, and whether gene was also differentially expressed in Orsay infected *rde-1* mutant data set (Chen et al., 2017). **B)** Overlap between genes significantly upregulated by RNA1(wt) in the *drh-1(+)* background with canonical IPR genes and between genes significantly upregulated by RNA1(wt) in the *drh-1(−)* background. **C)** Overlap between genes significantly upregulated by RNA1(wt) in the *drh-1(+)* background with canonical IPR genes and between genes significantly upregulated by Orsay infection from Chen et al 2017.

## Discussion

The field of innate immunity in *C. elegans* is only about 20 years old, and thus is relatively new compared to innate immunity research in other model hosts (Jiang and Wang, 2018; Kim and Ewbank, 2018; Leggewie and Schnettler, 2018; Xu and Cherry, 2014). While the transcriptional responses to diverse pathogens have been described in *C. elegans*, the pattern recognition receptors that activate these responses have remained mostly unclear, with the exception of a G-protein-coupled receptor required to sense ligands induced by both wounding and a fungal infection that penetrates the epidermis (Kim and Ewbank, 2018; Zugasti et al., 2014). Here we show that one of the only pattern recognition receptors conserved between mammals and *C. elegans*, DRH-1/RIG-I, appears to sense viral intermediates to trigger the IPR transcriptional program in the intestine, which is associated with increased defense against the Orsay virus and other intracellular pathogens (Reddy et al., 2019). Importantly, the role of *drh-1* in mediating the *C. elegans* transcriptional response to Orsay virus is separable from its role in antiviral RNAi, as neither of the two known direct binding partners of DRH-1, RDE-4 or DCR-1, nor other RNAi pathway components we tested, were required for IPR activation. Because the RDRP activity from the Orsay RNA1 segment appears to be required for activation, our results suggest that a viral replication intermediate, such as dsRNA or 5’ triphosphate RNA, is the ligand that binds the DRH-1 receptor to activate the IPR (Figures 4–6). This pathogen ligand/pattern-recognition receptor pair is one of the few that have paired together so far in *C. elegans*. Of note we found that microsporidia induction of the IPR is independent of *drh-1*, indicating that there are separate receptors yet to be identified that sense infection with these fungal-related pathogens. It is possible that microsporidia are sensed by the proteotoxic stress they cause during infection, although given the large number of effectors secreted by microsporidia (Reinke et al., 2017), there may be also be receptors that sense specific microsporidia ligands to trigger the IPR.

What happens downstream of DRH-1 activation to induce IPR gene expression? *C. elegans* does not have clear homologs of MAVS, IRF3, or NF-κB, which activate the transcriptional program downstream of RIG-I-like receptors in mammals (Gebhardt et al., 2017). Therefore, the signaling components that mediate this response in *C. elegans* are likely to be different. So far the only other host factors shown to regulate activation of the IPR are the antagonistic paralogs *pals-22* and *pals-25*, which repress and activate the IPR respectively (Reddy et al., 2019). *drh-1* appears to act in parallel to both *pals-22* and external triggers of the IPR like microsporidian infection and proteotoxic stress. These findings indicate that *drh-1* is just one of multiple inputs to the IPR (Figure 7). While constitutive activation of IPR gene expression in *pals-22* mutants indicates that this transcriptional program provides anti-viral defense (Reddy et al., 2019), the contribution of *drh-1*-mediated induction of this program is less clear. In previous studies it appeared that *drh-1* might have a role independent of RNAi in anti-viral defense, as loss of both *drh-1* and RNAi factors together had greater susceptibility to infection compared to loss of the RNAi factors alone (Ashe et al., 2013). We also observed a trend towards higher accumulation of Orsay RNA1 transcripts in the *rde-1(−); drh-1(−)* double mutant background compared to the *rde-1(−)* single mutant background (Figure 5B), which would be consistent with this model. Indeed, *drh-1* does not appear to be required for antiviral siRNA biogenesis, but rather appears to regulate which regions of the viral RNA are converted into small interfering RNAs (Coffman et al., 2017). It is not currently possible to completely separate the effects of *drh-1* activity on RNAi from its effects on IPR activation, but it is possible that some of the viral susceptibility in *drh-1* mutants that has been attributed to defects in antiviral RNAi could instead be due to defects in IPR activation.

**Figure 7:**
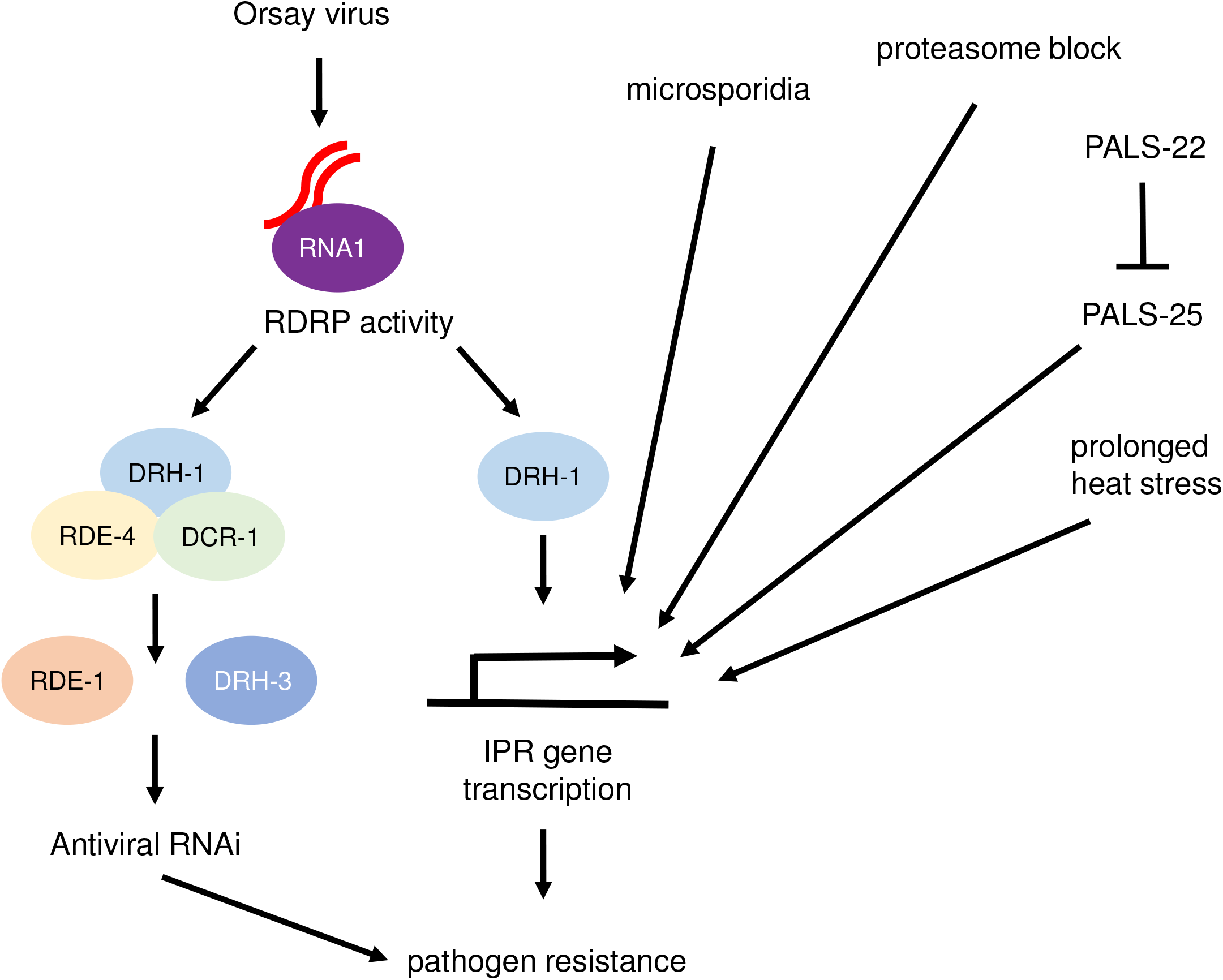
Model of DRH-1 mediated IPR activation by Orsay virus. Orsay virus infection in *C. elegans* is detected by DRH-1 recognition of viral replication intermediates produced by Orsay RNA-dependent RNA polymerase activity. DRH-1 signals downstream to activate the transcription of the protective IPR response. In parallel, DRH-1 also participates in the production of antiviral RNAi. The IPR can also be triggered by microsporidian infection, prolonged heat stress, proteasome blockade, and mutations in *pals-22*. Unlike viral infection, these triggers do not require DRH-1.

In mammals, a prominent output of RIG-I signaling is transcriptional upregulation of interferon genes that encode ligands to activate interferon receptors and downstream signaling. Because *C. elegans* lacks interferon orthologs, it will be interesting to determine the nature of IPR effectors and how they regulate resistance. Of note, *C. elegans* does have homologs of the STAT transcription factors, which act downstream of the interferon receptor in mammals to activate interferon-stimulated genes and promote anti-viral defense (Dierking et al., 2011; Schneider et al., 2014; Wang and Levy, 2006). Surprisingly, the *sta-1* STAT transcription factor appears to repress expression of viral-response genes, and *sta-1* mutants are resistant to infection (Tanguy et al., 2017). This result highlights another difference between mammals and *C. elegans* in terms of the signaling events downstream of RIG-I recognition of viral infection.

One intriguing theme in common between RIG-I/IPR signaling in *C. elegans* and RIG-I/interferon response in mammals is that defects in the proteasome are associated with both responses. Previous work in *C. elegans* demonstrated that either genetic or pharmacological block of the proteasome will activate IPR gene expression, in a manner independent of canonical proteostasis factors like the SKN-1/Nrf2 transcription factor (Bakowski et al., 2014; Reddy et al., 2019). In humans, mutations in the proteasome, as well as gain-of-function mutations in RIG-I-like receptors, can lead to inappropriate activation of type-I interferon responses and autoimmunity (Brehm et al., 2015). These defects are part of a group of diseases called interferonopathies, which are associated with upregulation of interferon, although it is controversial whether interferon is causal for these diseases (Kretschmer and Lee-Kirsch, 2017; Uggenti et al., 2019). Therefore, it is possible that mammalian RIG-I triggers an interferon-independent output similar to the IPR that promotes both immunity as well as damaging inflammation. In light of this idea, it is interesting to note that constitutive activation of the IPR in *pals-22* mutants is associated with increased immunity but also fitness defects such as shortened lifespan and premature aging (Reddy et al., 2017). Therefore, further analysis of the regulation and outputs of the IPR may shed light on the interplay between viral infection, immunity and the negative consequences of hyper-activation of immune responses.

## Acknowledgements

The authors would like to thank R. Underwood, A. Birmingham, and K. Fisch for assistance with RNA-seq data analysis, and V. Lazetic, E. Tecle, S. Gang, I. Sfarcic, and T. Bui for suggestions on the manuscript. This work was supported by NIH under R01 AG052622 and GM114139 to ERT, and R21 AI133291 to DW. JNS was supported by NIGMS/NIH award K12GM068524. The project described was partially supported by the National Institutes of Health, Grant UL1TR001442 of CTSA. The content is solely the responsibility of the authors and does not necessarily represent the official views of the NIH. Some strains were provided by the CGC, which is funded by NIH Office of Research Infrastructure Programs (P40 OD010440).

---

**Supplemental table 1** Results of differential expression analyses for all RNA-seq comparisons performed in this study.

**Supplemental table 2** List of gene set overlaps from Figure 6a-b.

**Supplemental table 3** Comparison between genes upregulated in *rde-1* mutant background by Orsay virus infection (Chen et al., 2017) and gene expression changes in Orsay RNA1(wt) vs Orsay RNA1(mt) in *rde-1* mutant background (this study).

**Supplemental table 4** Normalized gene counts for all RNA-seq replicates from this study.

## Materials and Methods

### *C. elegans* strains and maintenance

*C. elegans* were maintained on nematode growth media (NGM) plates seeded with OP50 *E. coli* as previously described (Brenner, 1974). Worms were maintained at 20°C unless otherwise noted. See Table 1 for list of all *C. elegans* strains used in this study.

**Table 1.** List of all *C. elegans* strains used in this study.

| Strain designation | Genotype |
| --- | --- |
| ERT054 | *jyIs8[pals-5p::GFP, myo-2p::mCherry] X* |
| ERT712 | *drh-1(ok3495) IV; jyIs8[pals-5p::GFP, myo-2p::mCherry] X* |
| ERT286 | *pals-22(jy1) III; jyIs8[pals-5p::GFP, myo-2p::mCherry] X* |
| ERT782 | *pals-22(jy1) III; drh-1(ok3495) IV; jyIs8[pals-5p::GFP; myo-2p::mCherry] X* |
| N2 | N2 |
| WM27 | *rde-1(ne219) V* |
| WM49 | *rde-4(ne301) III* |
| RB2519 | *drh-1(ok3495) IV* |
| RB1024 | *drh-2(ok951) IV* |
| WM206 | *drh-3(ne4253) I* |
| ERT767 | *virEx23[HSP::OrsayRNA1; myo-2p::YFP]* |
| ERT768 | *virEx26[HSP::OrsayRNA1 D601A; myo-2p::YFP]* |
| ERT695 | *jyIs8[pals-5p::GFP; myo-2p::mCherry] X; virEx23[HSP::OrsayRNA1; Pmyo-2::YFP]* |
| ERT697 | *jyIs8[pals-5p::GFP; myo-2p::mCherry] X; virEx26[HSP::OrsayRNA1 D601A; Pmyo-2::YFP]* |
| ERT696 | *drh-1(ok3495) IV; jyIs8[pals-5p::GFP; myo-2p::mCherry] X; virEx23[HSP::OrsayRNA1; myo-2p::YFP]* |
| ERT698 | *drh-1(ok3495) IV; jyIs8[pals-5p::GFP; myo-2p::mCherry] X; virEx26[HSP::OrsayRNA1 D601A; myo-2p::YFP]* |
| WUM51 | *rde-1(ne219) V; jyIs8[pals-5p::GFP; myo-2p::mCherry] X; virEx23[HSP::OrsayRNA1; myo-2p::YFP]* |
| ERT808 | *rde-4(ne301) III; jyIs8[pals-5p::GFP; myo-2p::mCherry] X; virEx23[HSP::OrsayRNA1; myo-2p::YFP]* |
| ERT802 | *drh-3(ne4253) I; jyIs8[pals-5p::GFP; myo-2p::mCherry] X; virEx23[HSP::OrsayRNA1; myo-2p::YFP]* |
| ERT765 | *drh-1(ok3495) IV; virEx23[HSP::OrsayRNA1; myo-2p::YFP]* |
| ERT766 | *drh-1(ok3495) IV; virEx26[HSP::OrsayRNA1 D601A; myo-2p::YFP]* |
| ERT805 | *rde-1(ne219) V; virEx23[HSP::OrsayRNA1; myo-2p::YFP]* |
| ERT806 | *rde-1(ne219) V; virEx26[HSP::OrsayRNA1 D601A; myo-2p::YFP]* |
| ERT803 | *drh-1(ok3495) IV; rde-1(ne219) V; virEx23[HSP::OrsayRNA1; myo-2p::YFP]* |
| ERT804 | *drh-1(ok3495) IV; rde-1(ne219) V; virEx26[HSP::OrsayRNA1 D601A; myo-2p::YFP]* |
| ERT780 | *drh-1(jy110) IV; jyIs8[pals-5p::GFP; myo-2p::mCherry] X* |
| ERT709 | *drh-1(jy110) IV* |

### CRISPR deletion of *drh-1* locus

To generate a mutant with a deletion of the entire *drh-1* genomic locus we used the CRISPR co-conversion strategy (Arribere et al., 2014). Two crRNAs targeting either side of the *drh-1* genomic locus were designed using Chopchop (https://chopchop.cbu.uib.no/): GCGTCTCTCTACTAATACAC and GGTTTTGGTCATCTTGATGT. *drh-1* crRNAs, *dpy-10* crRNA, and tracrRNA were obtained from IDT and resuspended in IDT nuclease-free duplex buffer. CRISPR injection mix was constructed containing 0.5 μl 100μM *dpy-10* crRNA, 0.5 μl 100 μM *drh-1* crRNAs, 2.5 μl 100 μM tracrRNA tracrRNA, and 3.5 μl Cas9 (QB3 MacroLab). The Cas9 mixture was then injected into the gonad of approximately 30 young adult N2 worms. Injected worms were singled onto individual NGM OP50 plates and incubated at 20°C for 3-4 days. Plates with a large number of F1 progeny showing the Dpy phenotype were identified, and >100 individual Dpy+ F1 progeny were picked to individual plates. Once F1 progeny had laid eggs, they were lysed and genotyped for the deletion of the *drh-1* locus using primers deletion external forward: CTCGTCACCAGTGCGAAATA and deletion internal reverse: CCAACCGCAATTCCAACATC. Presence of the complete deletion was confirmed with Sanger sequencing, and the allele was designated *drh-1(jy110)*. The *dpy-10* mutation was crossed out of the *drh-1(jy110)* background, and *drh-1(jy110)* mutants were backcrossed three times to N2 prior to use in experiments.

### Orsay virus filtrate preparation

Orsay virus filtrates were prepared as previously described (Félix et al., 2011). Briefly, infected *rde-1(ne219)* worms were grown on standard NGM OP50 plates until just starved, then washed off plates using M9 buffer and disrupted with silicon beads. The homogenate was then filtered through a 0.22 μm filter and aliquots flash frozen in liquid nitrogen.

### Orsay virus infection

For L1 infection (Figure 1a), adults were bleached to obtain synchronized L1 larvae, which were then mixed with 10X concentrated OP50 *E. coli* and 1:50 diluted Orsay virus filtrate. Then, 500 μL total volume of L1/food/virus mix was plated onto 6 cm unseeded NGM plates and dried in a laminar flow hood. Infected worms were then incubated at 20°C until collection.

For L2 infection (Figure 1b-f), bleached L1 larvae were plated on 6cm NGM plates containing a lawn of OP50 *E. coli* and incubated at 20°C overnight prior to infection. Orsay virus filtrate was then diluted in M9 at a ratio of 1:50. 300 μl of 1:50 diluted filtrate was top-plated onto the plates, which were then dried in a laminar flow hood. Infected worms were incubated at 20°C for 24h prior to collection.

For L4 infection (Figure 3), 10 μL of Orsay filtrate was mixed with 300 μL of 10X concentrated OP50 *E. coli* and 190 μL M9, and 500 μL of this mixture was seeded onto 6 cm NGM plates and dried in a laminar flow hood. L4 stage worms (bleached L1 larvae plated on standard 6 cm OP50 plates and incubated at 20°C for 2 days) were washed off plates using M9 + 0.1% Triton X-100 (TX-100), counted, and re-plated onto plates seeded with Orsay filtrate mix. Infected worms were incubated at 20°C for the indicated amount of time prior to collection.

### *pals-5p∷GFP* quantification

*pals-5p∷GFP* quantification by worm sorter (Figure 1, Figure 2a-h)

Worms were washed off plates into microcentrifuge tubes using M9 + 0.1% TX-100, then washed 3X with M9 + 0.1% TX-100. Worms were concentrated into 150-200 μl, and transferred into 96 well cell culture plates. Worms were then analyzed using a COPAS Biosort (Union Biometrica) to record time of flight (TOF) and green fluorescence for each worm. GFP signal was normalized to worm size by dividing fluorescence by TOF for each worm.

*pals-5p∷GFP* quantification by ImageJ (Figure 2i, Figure 3a-b)

Worms were collected and washed as described above, then paralyzed by adding 2 μL 5M Sodium Azide. Worms were then mounted on 2% agarose pads on glass slides, sealed with coverslip, and imaged using a Zeiss AxioImager M1 upright fluorescent microscope with a 10X objective. Signal was collected for DIC, YFP (co-injection marker for heat shock Orsay RNA1 expression array), GFP (*pals-5p∷GFP*), and mCherry (*pals-5p∷GFP* co-injection maker). Identical exposure times were used for all images within an experiment. GFP signal in the intestine of individual worms was quantified using ImageJ (https://imagej.nih.gov/ij/). Mean grey values for each intestine were normalized by subtracting mean grey value from the image background.

### *N. parisii* infection

*N. parisii* ERTm1 spore filtrate was prepared as previously described (Estes et al., 2011). 1200 bleached L1 larvae were combined with 500,000 ERTm1 spores and 150 μl 10X concentrated OP50 *E. coli* and M9 buffer to a total volume of 300 μl, then seeded onto 6 cm NGM plates and dried in a laminar flow hood. ERTm1 infected worms were incubated at 25°C for 30 h prior to collection.

### *N. parisii* pathogen load

N. *parisii* pathogen load was assessed by FISH staining as previously described (Reddy et al., 2019). Briefly, infected worms were fixed in 4% paraformaldehyde, then hybridized with a CalFluor610-tagged FISH probe (Biosearch) specific for *N. parisii* rRNA. FISH stained worms were analyzed using a COPAS Biosort (Union Biometrica) to record TOF and red fluorescence for each individual worm. Signal was normalized to worm size by dividing red fluorescence over TOF for each worm.

### Bortezomib treatment

Bleached L1 larvae were plated onto 6 cm NGM plates containing a lawn of OP50 *E. coli* and incubated at 20°C overnight. A 10 μM stock solution of bortezomib (Selleck Chemicals) resuspended in DMSO was mixed with M9 and top-plated onto plates for a final concentration of 2.5 μM bortezomib per plate. Control plates were top-plated with an equal amount of DMSO in M9. Plates were then incubated at 20°C for 24h prior to analysis with the COPAS Biosort (Union Biometrica).

### Prolonged heat stress

Bleached L1 larvae were plated onto 6 cm NGM plates containing a lawn of OP50 *E. coli* and incubated at 20°C overnight. Experimental plates were then incubated at 28°C for 24 h, while control plates remained at 20°C. Worms were analyzed after 24 h using the COPAS Biosort (Union Biometrica).

### qRT-PCR

Worms were washed off plates and washed with M9, then concentrated into <50 μl. Worms were homogenized in TRI Reagent (Molecular Research Center, Inc.) and frozen at −80°C. RNA was extracted using TRI Reagent and 1–bromo–3–chloropropane (BCP) (Molecular Research Center, http://www.mrcgene.com) according to the manufacturer’s instructions. cDNA was prepared from total RNA using either SuperScript VILO (ThermoFisher) or iScript (Bio-Rad) cDNA synthesis kits. qRT-PCR was performed using iQ SYBR Green Supermix (Bio-Rad) on a CFX Connect Real Time System. Each experimental replicate was measured in technical duplicate. All gene expression was normalized to *snb-1* expression, which does not change upon conditions tested. For comparisons in strains containing the *Ex[HSP∷RNA1]* arrays, gene expression was additionally normalized to *yfp* expression to control for array mosaicism and differing numbers of array-containing animals in the test populations. The Pffafl method was used for quantifying gene expression changes (Pfaffl, 2001). For list of qRT-PCR primers, see Table 2.

**Table 2.**
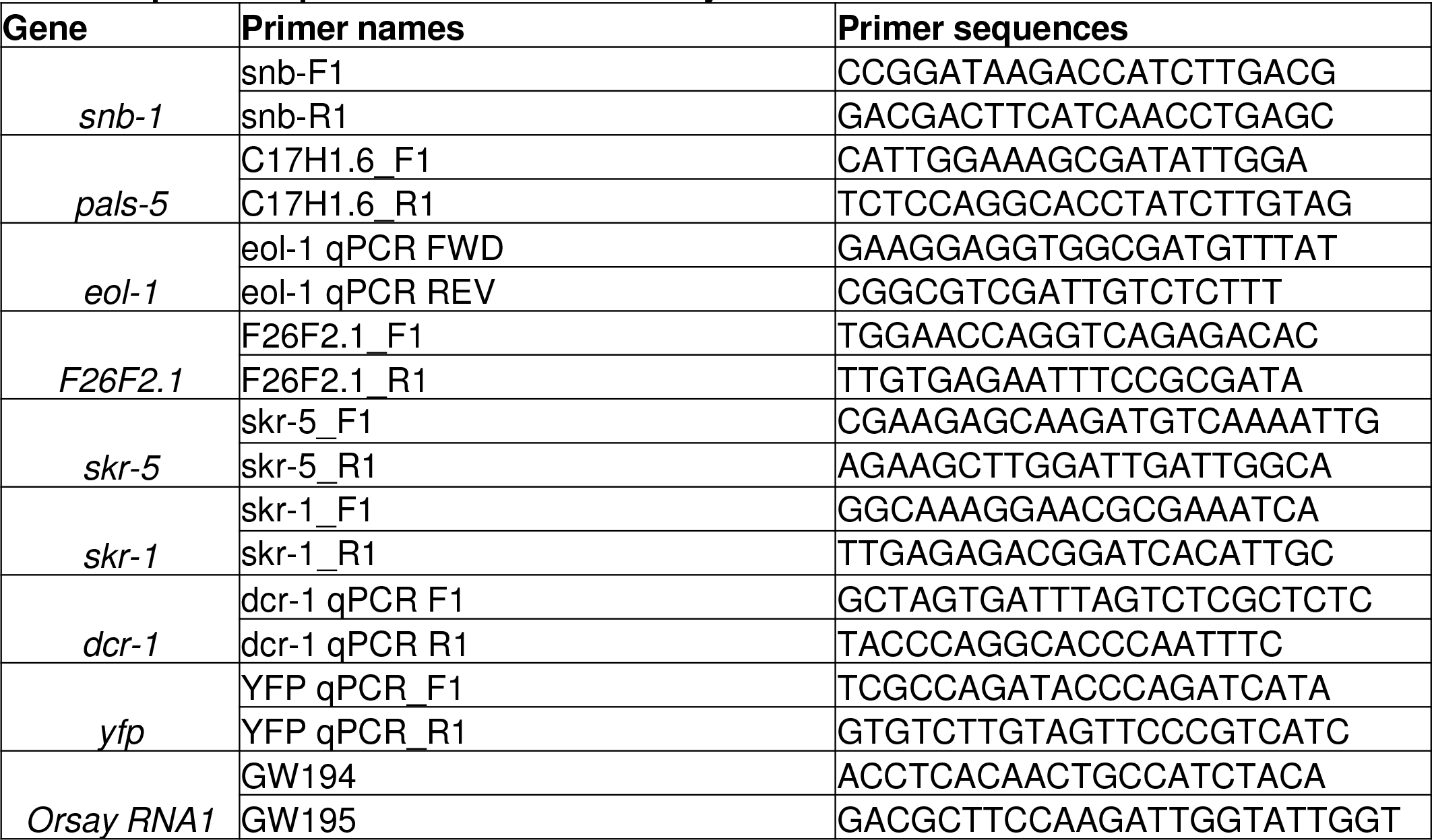
List of all qRT-PCR primers used in this study.

### *dcr-1* RNAi treatment

Overnight cultures of Ahringer library *dcr-1 E. coli* RNAi clone or RNAi vector control L4440 *E. coli*. clone in the HT115 bacterial strain were seeded onto 6 cm NGM plates supplemented with 5 mM IPTG and 1 mM carbenicillin and incubated at room temperature for 1 day. Synchronized L1 larvae obtained by bleaching were plated on RNAi *E. coli* lawns and grown for 48h at 20°C. 300 μl of 1:50 diluted Orsay virus filtrate was top-plated onto the plates, and plates dried in a laminar flow hood. Infected worms were incubated at 20°C for 24 h prior to collection

### HSP∷RNA1 heat shock induction

Bleached L1 larvae were plated on 6 cm NGM plates containing a lawn of OP50 *E. coli* and incubated at 20°C for 2 days (until L4 stage). At L4 stage, plates were heat shocked at 34°C for 2 h, then allowed to recover at 20°C for 6 h prior to collection.

### RNA-seq sample preparation and sequencing

Bleached L1 larvae were sorted using a COPAS Biosort (Union Biometrica) to obtain a population enriched for the *Ex[HSP∷RNA1(wt)]* or *Ex[HSP∷RNA1(mt)]* extrachromosomal array. Array-positive larvae were sorted to unseeded 10 cm NGM worm plates at a density of ~2000 worms/plate, and 1 ml 10X concentrated OP50 *E. coli* was added after sorting was complete. Plates were incubated at 20°C for 48 hours, until L4 stage. At L4 stage, plates were heat shocked at 34°C for 2h, then allowed to recover at 20°C for 6h prior to collection. Worms were washed off plates and washed 3X with M9, then concentrated into < 50 μl. Worms were then homogenized in 1mL TRI Reagent (Molecular Research Center, Inc.) and frozen at −80°C. RNA was extracted using TRI Reagent and 1–bromo–3–chloropropane (BCP) (Molecular Research Center, http://www.mrcgene.com) according to the manufacturer’s instructions, and additionally purified using the RNeasy clean-up kit with on-column DNase I digestion (Qiagen). RNA quality was assessed by TapeStation at the UC San Diego Institute for Genomic Medicine. Sequencing libraries were constructed using the Truseq stranded mRNA method (Illumina), and sequenced SR75 on an Illumina HiSeq4000 sequencer (Illumina).

### RNA-seq analysis

RNA-seq analysis was performed by the Center for Computational Biology and Bioinformatics at UC San Diego. Quality control of the raw fastq files was performed using the software tool FastQC (Andrews, 2010) v0.11.3. Sequencing reads were trimmed with Trimmomatic (Bolger et al., 2014) v0.36 and aligned to the *C. elegans* genome WBcel235 (Zerbino et al., 2018) using the STAR aligner (Dobin et al., 2013) v2.5.3a. Read quantification was performed with RSEM (Li and Dewey, 2011) v1.3.0 and the WBcel235 v96 annotation (Frankish et al., 2019). The R BioConductor packages edgeR (Robinson et al., 2010) and limma (Ritchie et al., 2015) were used to implement the limma-voom (Law et al., 2014) method for differential expression analysis. In brief, lowly expressed genes—those not having counts per million (cpm) ≥ 1 in at least 1 of the samples—were filtered out and then trimmed mean of M-values (TMM) (Robinson and Oshlack, 2010) normalization was applied. The experimental design was modeled upon genotype and treatment (~0 + genotype + treatment). The voom method was employed to model the mean-variance relationship in the log-cpm values, after which lmFit was used to fit per-gene linear models and empirical Bayes moderation was applied with the eBayes function. Significance was defined by using an adjusted p-value cut-off of 0.05 after multiple testing correction (Benjamini and Hochberg, 1995) using a moderated t-statistic in limma. For gene lists, Wormbase version WS270 was used to remove dead genes and update gene names.

### Statistics

Statistics were performed using Prism 7.

